# Degradation of organic matter by ecologically distinct microbiomes uniformly promotes plant growth

**DOI:** 10.1101/2024.10.27.620506

**Authors:** Bradie Lee, Devin Coleman-Derr, John P. Vogel

## Abstract

In this study, we explore the relationship between ecologically distinct soil microbiomes and their role in plant growth promotion through organic matter (OM) decomposition. Using a controlled experimental system, we tested whether living microbiomes from geographically and ecologically distinct environments impact plant biomass when provided with OM in the form of dried, ground leaves. Results showed a consistent threefold increase in plant biomass when a living microbiome and organic matter were present, regardless of the microbiome’s origin, which included agricultural fields, desert soil, and pine-oak forest soil, municipal compost, and the microbiome on unautoclaved organic matter. Bacterial community profiling based on 16S amplicon sequencing revealed genera, e.g. *Massilia*, that were significantly associated with decomposition and subsequent plant growth promotion. This suggests a conserved functional capacity for organic matter decomposition across diverse microbiomes, likely due to evolutionary pressures to efficiently break down plant material for nutrient acquisition. The study provides a framework for further investigation into microbial consortia that enhance plant growth via decomposition, offering a robust experimental system to identify microbes and microbial processes that could be harnessed to improve nutrient uptake from organic inputs like cover crops.

## INTRODUCTION

The interaction between plants and the soil microbiome is a highly complex and often mutually beneficial relationship. Plants release small organic compounds, such as sugars and amino acids through exudation from their roots into the soil (Sasse, Martinoia, and Northen 2017). Plants also provide more complex organic matter (OM) to the soil when they die or shed tissues (e.g. sloughed root cap cells and fallen leaves) (Cotrufo et al. 2013). Collectively, these organic materials serve as food sources for microbial communities, and in turn these microbes, including bacteria, perform various essential functions including, but not limited to, nutrient cycling that supports plant growth (Das et al. 2022). These processes release nutrients locked in complex molecules into simpler forms that plants can readily absorb through their root systems. This symbiotic relationship is vital for ecosystem function, as plants heavily rely on microbial decomposition to access nutrients that would otherwise remain locked in organic matter (Das et al. 2022; Lambers et al. 2008). Conversely, the microbiome depends on plants for the carbon and energy needed to sustain its activities, creating a tight feedback loop where each party plays a critical role in the other’s survival (Zhalnina et al. 2018; L. Li et al. 2024). Without this intricate cooperation, plants would struggle to thrive, particularly in nutrient-poor environments, and the diversity and function of soil microbial communities would be compromised. However, relatively little is known about how bacterial communities function in the context of organic matter cycling, and this represents a key knowledge gap for future efforts to manipulate microbiomes for increased nutrient uptake by plants in agricultural settings.

Organic matter decomposition is a complex process that plays a crucial role in nutrient cycling and ecosystem functioning. While both biotic and abiotic factors contribute to decomposition, most decomposition is mediated by microbial processes. Microbial decomposition can be influenced by a variety of abiotic factors, such as temperature, water content, and pH (Raza et al. 2023); microbiome composition and function have also been shown to vary along these clines (Bahram et al. 2018). Microbes such as bacteria and fungi break down organic materials through enzymatic activity, converting complex compounds into simpler, bioavailable forms that plants can acquire. For example, bacteria excrete enzymes such as proteases that depolymerize complex molecules, resulting in forms of nitrogen that can be acquired by the plant (Sieradzki et al. 2023). Similarly, fungi can break down phytate, a complex molecule that plants use to store phosphorus in their cells but cannot absorb from the environment (Adedayo and Babalola 2023). These and other mechanisms can be found in a wide variety of organisms involved in decomposition. While microbial communities are known to drive much of the decomposition process, little is understood about how the ecological origin of these communities influences their ability to decompose plant matter or influence plant growth via decomposition.

Common farming practices (e.g. cover cropping and compost amendments) suggest that the decomposition of organic matter promotes plant growth (e.g. (Tonitto, David, and Drinkwater 2006; Arsjad 1963; Wagger 1989), however, few studies have elucidated the complex mechanisms governing the diverse interactions between species in these systems. Synthetic bacterial communities have been shown to increase plant growth via decomposition of complex carbon molecules (Zhang et al. 2019; Zhou et al. 2024), and arbuscular mycorrhizal fungus have been shown to act synergistically with the bulk microbiome to promote nutrient uptake from leaf litter (Hestrin et al. 2019). Microbiome origin has been shown to affect leaf litter decomposition, with some microbiomes showing higher capacity to degrade their native litter type (Ayres et al. 2009). Taken together this suggests that origin of a microbiome origin may impact its ability to promote plant growth through the decomposition of particular organic materials due to specific community dynamics. However, to our knowledge, no studies have demonstrated whether diverse, ecologically divergent soil microbiomes vary in capacity to promote plant growth via decomposition of a specific source of OM. In this study we tested the ability of divergent microbiomes to promote plant growth via decomposition of dried grass leaves. Surprisingly, we observed similar growth promotion irrespective of microbiome origin. Additionally, we highlight specific microbes that are shared by all the studied microbiomes that may contribute to plant growth promotion

## RESULTS

### 2.1 Living microbiomes increase plant growth in the experimental system

To determine the role of a living microbiome in plant growth promotion from the addition of organic matter (OM), we performed an experiment in which plants were grown either in the presence or absence of soil tea (microbiome source) and additionally in the presence or absence of OM (dried grass leaves). Both autoclaved and unautoclaved combinations of each treatment were also included (Figure 1A). The treatments not receiving a combination of a living microbiome (from soil tea or unautoclaved organic matter) and organic matter served as negative controls, and no growth promotion was expected. Plants were harvested after 4 weeks of growth. The shoots were dried to determine above ground biomass, and the roots were processed for microbiome characterization. Pairwise comparisons of dry biomass yield revealed statistically significant impacts of all the treatment factors (Figure 1B), including OM addition, inoculation type, and sterility. We observed an approximately 3-fold increase in biomass in plants that received both OM and a living microbiome compared to all of the sterile treatment combinations as well as the treatment receiving the unautoclaved soil tea only. The only significant interaction term in the ANOVA test was the interaction between the presence or absence of a living microbiome and OM (p < 0.001). Collectively, these data confirm that in this experimental system, the presence of living microbes is necessary for plants to experience growth promotion from the presence of OM within the media.

**Figure 1.**
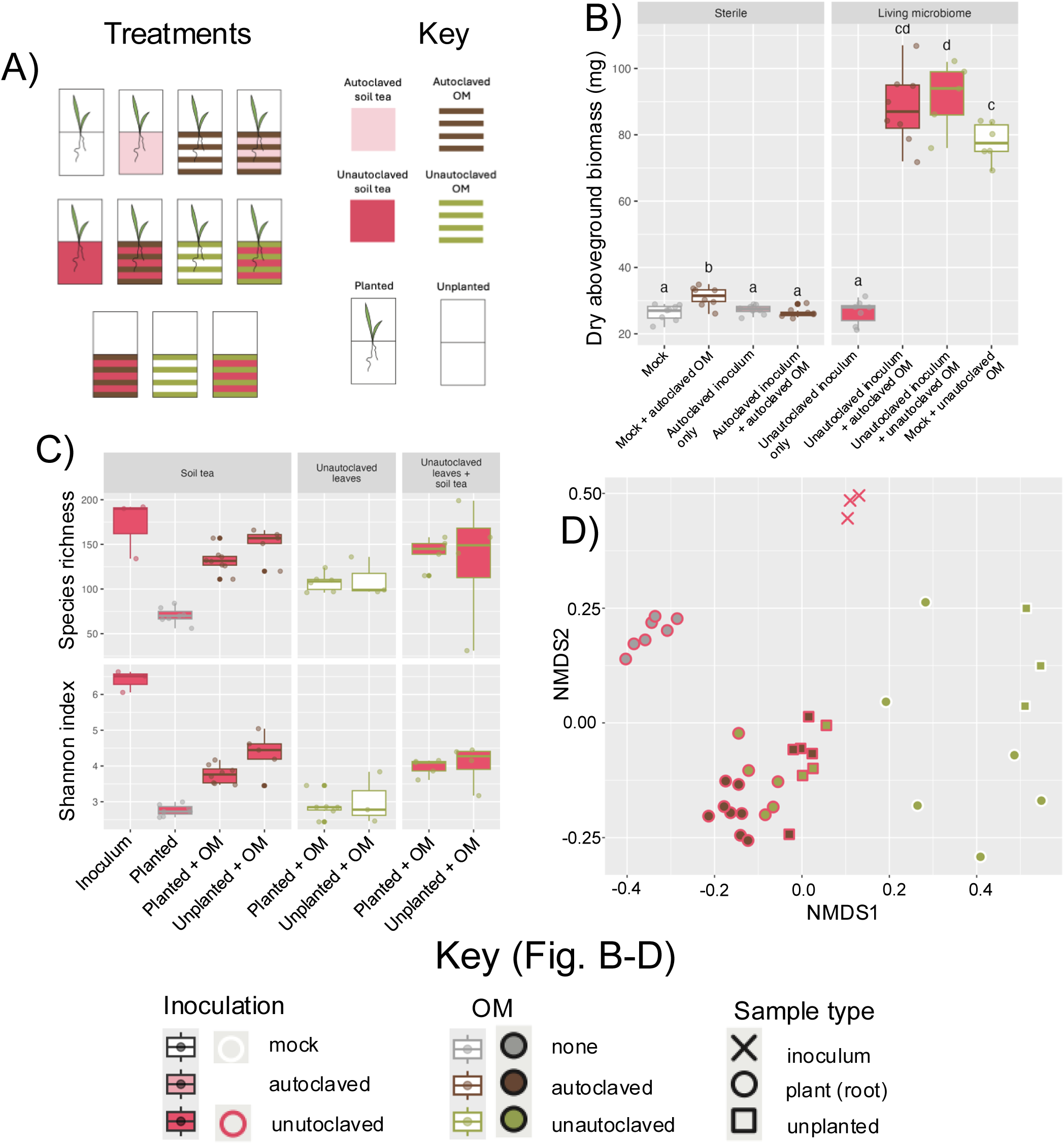
Experiment showing living microbiomes are responsible for plant growth promotion phenotypes due to OM, displaying experimental design, phenotypes, and microbiome structures associated with each treatment type. A) Experimental design showing all combinations of experimental treatments utilized in the phenotype and microbiome analyses. B) Total dry aboveground biomass in mg for plants in each experimental microcosm. After filtering samples of mitochondrial and chloroplast reads, retaining those with above 5000 reads in samples receiving living microbiomes, C) Species richness and Shannon diversity in samples that received a living microbiome, and D) NMDS analysis shows axes 1 and 2 of a Bray-Curtis dissimilarity matrix calculated on all samples that received a living microbiome.

### 2.2 Experimental conditions significantly impact microbiome dynamics

Prior research has shown that the presence of organic material can play a role in determining soil community composition (Cesarano et al. 2017; Whitman et al. 2016; Testen and Miller 2018). To determine whether the addition of OM treatment in our system impacted the microbiome diversity or composition within the root microbiome, DNA extraction and 16S amplicon sequencing was performed on the root samples. As expected, most samples that did not receive a living microbiome did not produce a sufficient number of 16S reads to analyze due to the presumed sterility of these treatments. As a result, these samples were excluded from the remaining 16S analyses. Additionally, we were unable to extract enough DNA to analyze from the inoculated, unplanted samples without OM suggesting that, in this experimental system, the microbes added from the inoculum can only persist when organic substrates are provided, either through the living plant roots and their exudates or added OM.

To explore how microbial diversity in this system was impacted by OM and inoculum treatments, Shannon’s diversity and species richness were determined (Figure 1C). ANOVA tests showed that while OM was a significant predictor of Shannon diversity (p < 0.0001) and richness (p < 0.0001), other factors were not. As anticipated, Tukey test comparisons of the inoculum to all other experimental samples indicate that the Shannon diversity dropped during the experiment compared to the starting point (p < 0.0001 for all comparisons to the inoculum). The inoculum had significantly more unique ASVs than the planted samples with only an inoculum or uninoculated, planted samples with unautoclaved OM (Tukey test, p < 0.05 for all comparisons). Additionally, we observed that inoculated plants not given OM had relatively low diversity (planted, no OM added) compared to the others, with the exception of the samples with unautoclaved OM and no soil tea (p < 0.0001). Interestingly, microboxes inoculated with living microbes and supplemented with organic matter, both those with plants in them and those without, had relatively high diversity and species richness; the only treatment group with higher diversity was the inoculum itself. The alpha diversity and richness metrics, in addition to the sample quality control, indicate that only a subset of microbes from the original inoculum persisted in all samples. The final community composition generally had fewer bacterial species, less even abundances, or both, and the amount and evenness that persisted depended on the starting microbiome as well as the substrates that were available to them.

Next, to understand whether the microbial communities in each sample type varied in composition we examined differences in beta diversity across our dataset. The samples clustered as distinct groups according to treatment in NMDS ordination plots, suggesting differences in composition by treatment group (Figure 1D). Congruent with the alpha diversity analysis, the inoculum itself was furthest from the rest of the samples, while the roots with no OM and a living inoculum were similarly tightly clustered and furthest from the other experimental samples. The samples with unautoclaved inoculum, regardless of the autoclave status of the OM, all clustered together compared to the rest of the treatments, but with some apparent separation across each treatment factor. Factors that appeared to influence axis 1 were the presence of OM and its autoclaved status, as well as whether the sample was planted or unplanted. PERMANOVA testing showed that the presence and autoclaved status of the OM (R^2^ = 0.31, p < 0.001), inoculation status (R^2^ = 0.13, p < 0.001), and sample type (R^2^ = 0.15, p < 0.001) significantly influenced community structure. An interaction between inoculation status and sample type was also observed (R^2^ = 0.05, p = 0.0147), but no other interaction terms were observed to be significant. The OM (none, autoclaved, or unautoclaved) had the greatest effect on the dataset, with approximately 31% of all variance being explained by this variable. Inoculation status also had a large effect on the dataset at approximately 13%, which was demonstrated by the mock inoculated samples with unautoclaved OM apparently clustering across axis 1. Collectively, these data suggest that both treatment factors, inoculum and OM supplementation clearly and significantly impacted the community assembly.

### 2.3 Distinct microbiomes uniformly promote plant growth in the presence of OM

Soil microbiomes are known to vary across many ecological factors (Si et al. 2021; Isobe et al. 2020; Naylor et al. 2017; Santos et al. 2021; Lauber et al. 2009) and we anticipated that different soils may provide more or less benefit to their plant hosts when organic matter was added. To determine whether the ecological history of different microbiomes influenced the plant growth promotion effect seen in the previous experiment, we utilized our experimental system to test the impact of microbiomes on plant growth from four distinct ecosystems: agricultural. soil, municipal compost, desert soil, and soil from a pine-oak forest (Figure 2A). In this second experiment, congruent results to the first experiment were generally obtained. For the aboveground biomass, all of the negative controls were significantly smaller than the treatments with OM and a living microbiome across all of the inoculation types; plants with OM and a living microbiome were approximately 3-fold larger than the negative controls (Figure 2B). An ANOVA test indicated a significant effect of both OM treatment and inoculation status as well as an interaction of the two (p < 0.001). As expected, post-hoc testing revealed that there was no significant difference between any of the samples that did not receive an unautoclaved inoculum and OM. Furthermore, a 3-fold increase in biomass was seen between the samples with living inoculum and organic matter and the negative controls without this specific combination, a result that is near-identical to that observed in the first experiment. However, contrary to our hypotheses, all samples that received a combination of unautoclaved inoculum and OM showed similar increases in biomass. Collectively, this indicates that the microbiomes adapted to diverse conditions tested here all possessed the capability of increasing plant growth via decomposition of OM.

**Figure 2.**
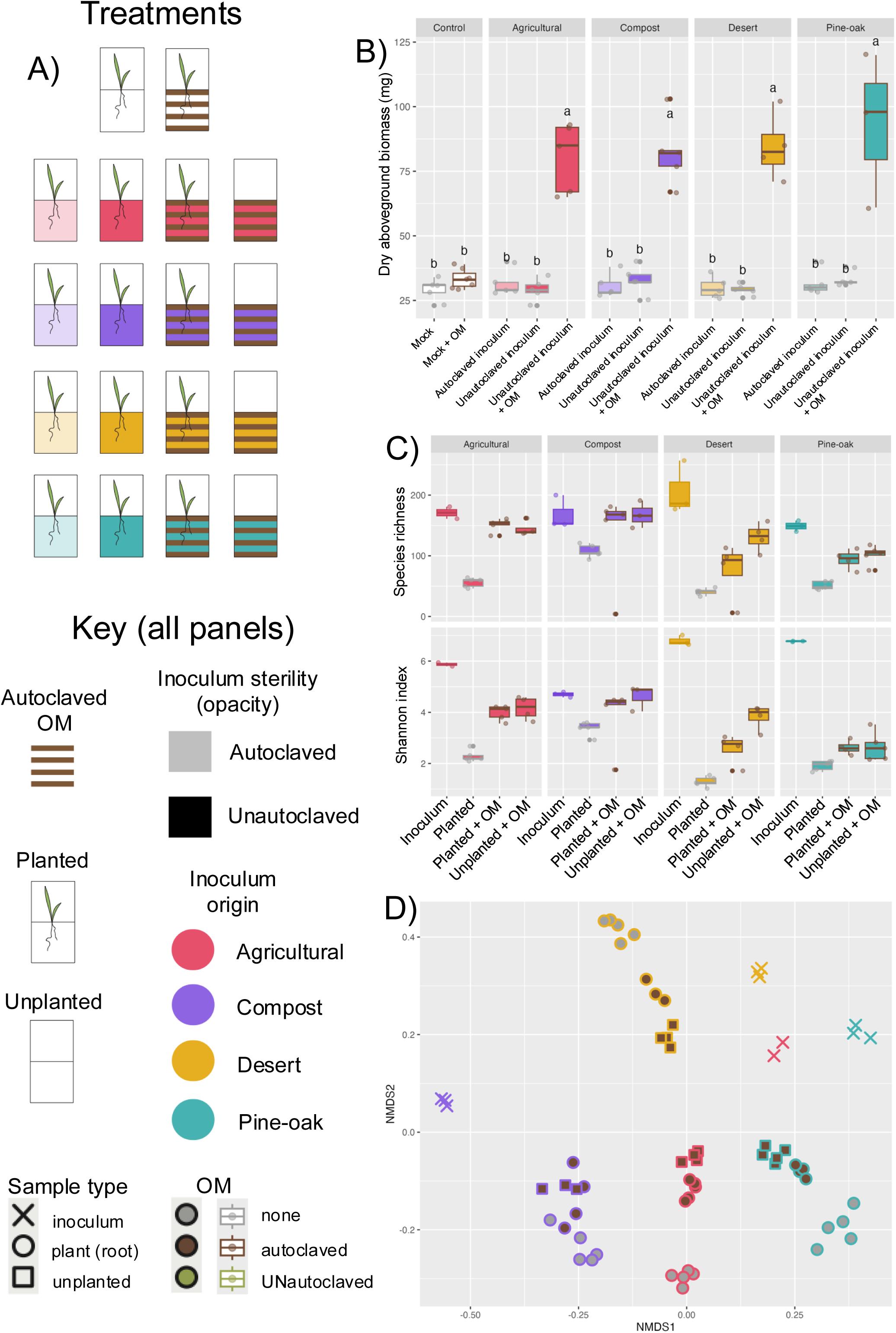
Microbiome comparisons experiment displaying experimental design, phenotypes, and microbiome structures associated with each treatment type. A) experimental design showing all combinations of experimental treatments utilized in the phenotype and microbiome analyses. B) aboveground biomass in grams as total weight in mg of all three plants in each experimental microcosm. After filtering samples of mitochondrial and chloroplast reads and retaining those with over 5000 reads, in samples receiving living microbiomes, C) Species richness and Shannon diversity index in samples that received a living microbiome above 5000 reads per sample, and D) shows axes 1 and 2 of a Bray-Curtis dissimilarity matrix calculated on all samples that received a living microbiome

### 2.4 Microbiome origin and experimental conditions interact to produce unique community dynamics

In the second experiment, we also explored the impact of the added soil tea on microbiome community composition Shannon diversity and species richness were broadly similar to our first experiment, but with distinct differences between inoculum types (Figure 2C). ANOVA showed OM alone did not significantly impact Shannon diversity, though its interaction with sample and inoculation types did (p < 0.0001). Tukey tests revealed the initial inoculum generally had higher Shannon diversity compared to experimental samples, except for compost treatments. Inoculum diversity also varied by origin. Samples within the same treatment showed differences based on inoculation type, with compost inoculum exhibiting the lowest Shannon diversity compared to other inocula, but unlike other inocula, retained diversity in the experimental conditions with the addition of OM. Notably, root samples without OM had significantly lower Shannon diversity, with desert samples showing the lowest. Species richness followed similar trends, with all factors and interactions significant (p < 0.0001), except OM alone. Tukey tests indicated higher species richness in inocula compared to the other treatment types, except compost (p < 0.05). Treatments with OM had higher richness than planted samples without OM, aligning with Shannon diversity metrics. These results suggest varying capacities for microbial persistence and functional redundancy based on inoculum origin.

Similar to the first experiment, the communities clustered distinctly according to treatment groups in NMDS ordination plots, with significant differences observed between sample types, the presence or absence of OM, and inoculum origin, as well as some of their interactions (Figure 2D). Notably, a stark difference between soils of different origins was observed, with distinct clustering patterns for each inoculum origin, suggesting that the source of the microbiome plays a crucial role in shaping community assembly and its response to OM, despite the fact that source did not impact plant growth. Interestingly and congruently, the interaction of inoculum origin and sample type had the second highest R2, and the interaction of inoculum origin and OM was higher than OM alone, demonstrating the unique trajectory of each inoculation type along the other treatment factors. PERMANOVA results showed that all tested factors, including inoculation status (R^2^ = 0.336, p < 0.00001), sample type (R^2^ = 0.11, p < 0.00001), presence or absence of OM (R^2^ = 0.044, p < 0.00001), and the interactions between inoculation and sample type (R^2^ = 0.23, p < 0.00001), as well as inoculation and OM presence (R^2^ = 0.098, p < 0.00001), significantly influenced community structure. This divergence in soil microbiome composition highlights the impact of ecological history of the starting microbiomes in determining microbiome structure in response to decomposition of OM.

### 2.5 Specific microbial taxa are associated with different inocula origins and plant growth phenotypes

We hypothesized that while the final microbiomes differed from one another in overall composition, there may be conserved shifts in specific bacterial lineages across all treatment groups that received OM. Congruent with the NMDS analyses, obvious differences in the abundance of various bacteria within each treatment type and between factors were apparent in the relative abundance plots (Figure 3A). ANCOM-BC2 was utilized to determine taxa with significantly different abundance between sample types in both experiments, with an emphasis on the second experiment. A variety of genera and ASVs were identified as being differentially abundant between experimental samples, with shared patterns in their enrichment (p <0.05 for all comparisons) (Figure 3B). Of particular interest were those that were enriched in response to the addition of OM (both with or without plants) when compared to samples with plants and no OM. These were considered to be decomposition-associated microbes because they were not enriched by the root but rather by the OM alone. Utilizing pheatmap to group similar patterns together, these appeared to represent the first 5 ASVs in figure 3B. In contrast, there were other ASVs that were enriched in the planted samples without OM, indicating an opposite pattern that could be considered root-associated. Additionally, there were significant differences in genera between inoculation types (figure S8). This indicated that there were likely specific genera or strain level differences between inoculations that drove the phenotypes seen in previous analyses.

**Figure 3.**
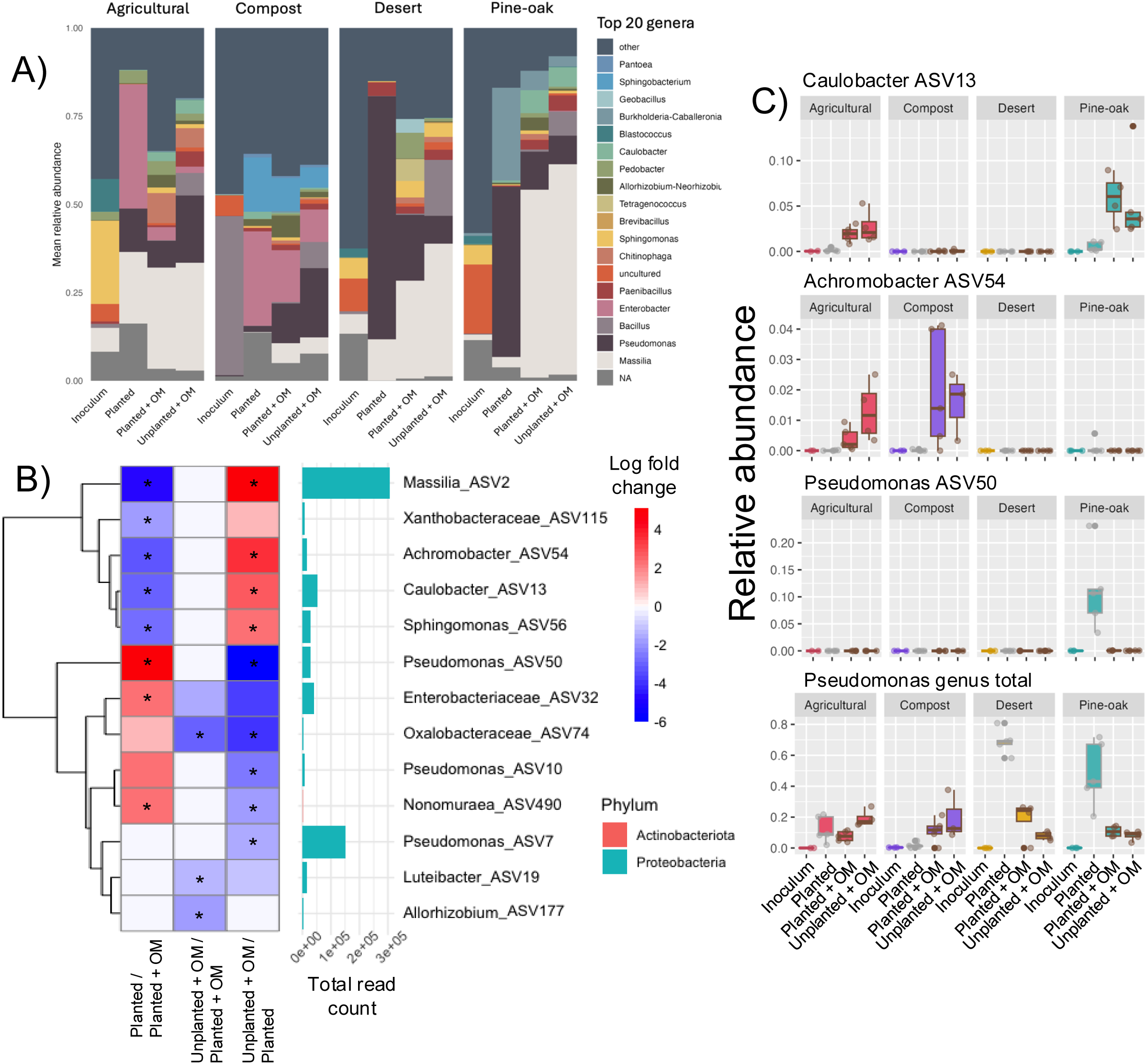
Microbiomes vary by inoculation and sample type. A) Relative abundance barplot displaying the mean abundance for the top 20 genera present in the samples that contained a living microbiome in Experiment B, faceted by inoculum type. Each bar is colored by the mean relative abundance per genera for all samples within each treatment type, with 3-5 samples per treatment. B) Heat map of log fold changes in relative abundance in ANCOM-BC2 analysis. All differential abundances displayed had one or more significant changes in abundance between sample type, with those passing the ANCOM-BC2 sensitivity test indicated by asterisks. Bars on right indicate total read count abundance, colored by phylum of each ASV. C) Relative abundance of select ASVs and corresponding genera that significantly varied between sample types in both the first and second experiment as well as inoculation type in the second experiment.

To further investigate how these significantly differentially abundant taxa varied between inoculation and sample types, we graphed the relative abundances of various significantly different taxa (Figure 3C). Between the first and second experiments, the only conserved pattern seen was in the Achromobacter and Caulobacter genera, which showed consistent enrichment in the presence of OM. The most abundant ASVs in each of these genera also showed this pattern of enrichment between experiments in the samples inoculated with the agricultural soil tea. These two genera were also significantly different between the inoculation types, where both were present in the agricultural samples, but the Caulobacter reads were present only in the pine-oak samples otherwise, and the Achromobacter reads were only present in the compost. Similar results were seen across a variety of other genera and ASVs (Figure 3B, S6-7), which showed a decomposition-associated enrichment pattern that varied in presence or abundance between inoculation types. These results show that many genera are associated with the decomposition process and are consequently associated with plant phenotype differences as well. Additionally, there may be taxonomically different microbial species or strains in ecologically distinct soils that could be responsible for the plant growth promotion.

Another taxa of interest due to its decomposition-associated enrichment in the second experiment is the Massilia genus; both the whole genus and select ASVs within this genus showed a decomposition-associated enrichment (Figure 4B, S6-7). The Massilia genus is one of the most common genera in the dataset, accounting for over 60% of reads in some samples, and was present in all inoculation types (Figure 4A-B, S1). Additionally, it was significantly depleted in the compost inoculum type compared to others (figure S8) and was significantly enriched in samples containing organic matter (Figure 3B). Interestingly, there was very little overlap between the individual *Massilia* ASVs found in samples containing each inoculation type despite similar patterns of abundance between sample types (Figure 4). This also held true for all ASVs in general (figure S3B). Interestingly, one of the most abundant Massilia ASVs (ASV2) was significantly enriched in samples with OM in 3 of the 4 inoculation types (Figure 4B). Massilia ASV3 was also found to be significantly enriched in OM-containing samples when ANCOM-BC2 analysis was performed on the agricultural samples only, and was only present in samples with this inoculation type. In the first experiment, *Massilia* as a genus was not found to have a decomposition-associated enrichment pattern (figure S4-5); indeed, when analyzing specific ASVs, some Massilia were actually found to be enriched in samples without OM. These individual ASVs were not the same ASVs that were found to be decomposition-associated in the second experiemnt, and differences in enrichment patterns across this genus suggest potential variation in affinity for OM presence within this lineage. These enrichments patterns may be due to the activity of genera that were unable to survive storage in the cold room, as the second experiment was conducted several months after the first. *Massilia* that are able to persist on living plants rather than degrading OM may not survive prolonged dormancy. However, because of its significant enrichment as a genus across all inoculation types in the second experiment, with unique ASVs enriched in different inoculation treatments, we hypothesize that some members of this genus may have some conserved function in the decomposition process within the root microbiome.

**Figure 4.**
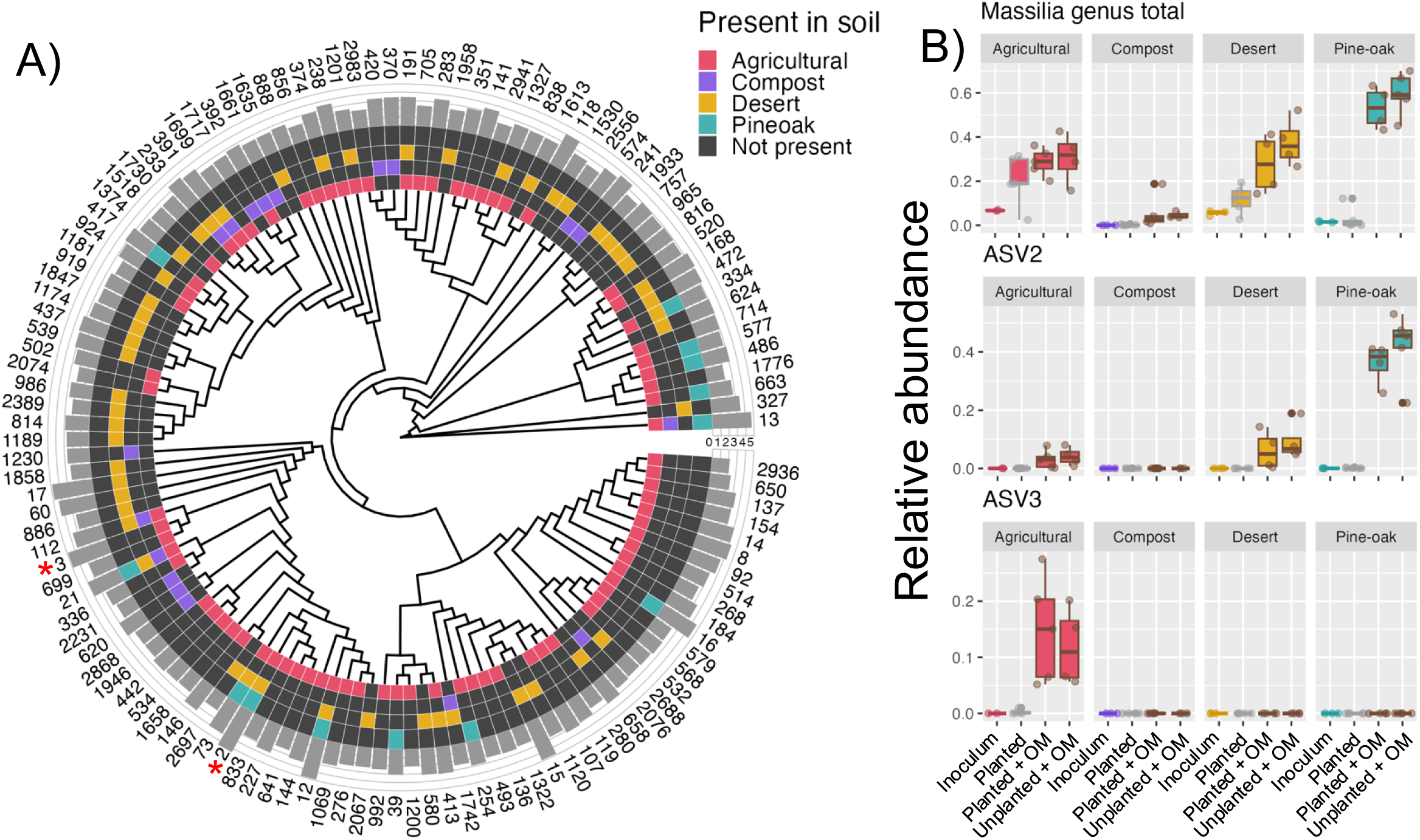
Massilia ASVs vary between experimental conditions. A) Phylogenetic tree of Massilia ASVs labeled by number corresponding to their total abundance in the combined dataset of all samples in both experiments that had above a total count of 100 reads in all combined experimental conditions in experiment B, rooted with Caulobacter ASV13. Colors represent presence or absence in each inoculum type, with corresponding bars representing the log scale total read abundance in the dataset. Asterisks indicate significantly differentially abundant ASVs displayed in B. B) Relative abundances (scale 0-1) of the Massilia genus in treatment conditions in experiment B as well as vidual ASVs 2 and 3.

In order to understand how similar the taxonomic composition of the microbiomes from each inoculation type were, we determined the presence/absence of specific ASVs in samples from each inoculation type. Despite near-identical phenotypes between inoculation types, many genera were significantly different in abundance (figure S8), and individual ASVs were generally found to be unique to each inoculation type. Out of the over 1100 unique ASVs with over 100 total reads identified in Experiment B, only 7 were found across all inoculation types (figure S3A), with one of these being classified as Massilia (Figure 4A). Massilia ASVs showed a similar pattern, with most of the individual ASVs being found only in one sample type (Figure 3A).

### 2.6 System robustness and reproducibility

Finally, to test the robustness of the experimental system, biomass yield and bacterial community composition from the two experiments described above, as well as biomass from a pilot experiment performed with an identical protocol, were compared to one another. All of the treatments with the living inoculum and OM present had no statistically significant differences. Some of the negative controls had significant differences, but all had a significantly smaller biomass compared to the treatments with living microbiomes and the presence of OM, with approximately 3-fold greater biomass (figure S9). Additionally, in a global PCoA analysis performed across all datasets, samples within treatment groups clustered most closely to analogous samples from other experiments (figure S2). Taken together, this global analysis indicates that although there was little overlap in the specific ASVs between experiments, conserved shifts in the community composition and plant phenotype are observable across differences in inoculum origin and experimental replicates.

## MATERIALS AND METHODS

A combination of treatments with the factors of soil/no soil, living/autoclaved soil, ground leaves or none, and planted/unplanted of the living soil treatments were utilized for this experiment. Control treatments contained only calcined clay and Hoagland’s solution, with the rest of the treatments described in further paragraphs.

### 3.1 Microbox Preparation

Microboxes (SacO2 Product ID O119/140+OD119/140) with #10 white filter lids were filled with 200 grams of calcined clay. Due to very high temperatures during manufacturing, calcined clay contains no organic matter except for trace amounts from dust contamination during handling. To each box in the leaf treatments, 1 gram of ground dried leaves was added. Hoagland’s solution was prepared as in (Novak et al. 2024), with slight modification (supplementary table 1) from stock solutions and titrated to pH 5.75 ± 0.05 using sulfuric acid. Each microbox received 110 mL of Hoagland’s solution and shaken to mix. Microboxes were autoclaved on a liquid/media cycle for 45 minutes. Trays containing the microboxes were wrapped with foil and taped shut prior to autoclaving. For the first experiment, 5-8 replicates were created for each experimental combination. For the second experiment, 5 replicates were made for each experimental combination.

### 3.2 Organic Matter Preparation

*Avena barbata* (wild oat) leaves were collected at LBNL before flowering between February and April in the spring of 2022 and 2023 using scissors to cut close to the soil line. The leaves were dried at 60°C in an oven until completely dried. The material was crushed by hand and passed through a 2 mm sieve and stored at room temperature for later use.

### 3.3 Soil collection

Soil samples were collected from their prospective sites by digging approximately 6 inches deep at each location, procuring one gallon ziplock bags of each soil, and storing the contents in plastic bags in a 4C cold room until use. Agricultural soil was collected from the UC Davis Century Plot with permission, at the coordinates 38.5370° N, 121.7869° W. The pine-oak soil was collected from lands managed by the California Bureau of Land Management at approximately 41.5925° N, 122.9269° W. The compost was collected from the Berkeley Marina free municipal compost pile at approximately 37.8722° N, 122.3159° W. The desert soil was collected from lands managed by the California Burea of Land Management at the coordinates 35.121111° N, 118.190000° W. The agricultural soil was harvested in June of 2022 while the rest were collected in the summer of 2023 between May and July.

### 3.4 Soil and Mock Inoculations

Soil tea was prepared by mixing 100 g of soil with 1000 mL of Milli-Q water using a stir bar for 30 minutes on medium-high speed. Large particles were allowed to settle for 5 minutes, and the liquid was poured off into separate flasks taking care not to disturb the settled soil. A 250 mL portion of the soil tea was autoclaved in a flask for 1 hour. Treatments were inoculated with 10 mL of soil tea (autoclaved or not depending on the treatment), and the remaining soil tea was capped and shaken between inoculations to ensure homgeneity. Control treatments received mock inoculations of 10 mL of vacuum sterilized Milli-Q water. The lids of filled microboxes were wrapped with Parafilm in a laminar flow hood. Inoculated microboxes were placed in a cold room overnight before planting.

### 3.5 Seed Sterilization and Planting

Seeds of *Brachypodium distachyon* accession Bd21-3 (Vogel and Hill 2008) were sterilized using a 70% ethanol rinse for 30 seconds with vortexing, followed by a 5 minute rinse in 5% sodium hypochlorite with vortexing. The seeds were then rinsed six times with sterilized water.

Sterilized seeds were placed on petri dishes containing 1/2 MS (Caisson, Smithfield, UT, USA; CatNo. MSP01-50LT) with 1.5% phytagel (Invitrogen, Waltham, MA, USA; Quant-iT dsDNA Assay Kit, CatNo. Q33120), MES at 0.5 g/L (Fisher Scientific, Waltham, MA, USA; CatNo. J61587.AP) and titrated to pH 5.7 with 1 M KOH. Petri plates were wrapped in foil and placed at 4°C for 5 days for stratification. After stratification, the plates were uncovered and placed vertically, with seeds oriented with embryos facing down, in a growth chamber to germinate.

Sterile treatments were planted first using autoclaved forceps. Healthy seedlings with similar root length were selected and placed in holes dug by the forceps, ensuring the root was pointing down, and then lightly covered with the growing medium. Three seedlings were planted in each microbox. Microbox lids were wrapped in Parafilm before removing them from the laminar flow hood. All microboxes, including unplanted ones, were randomly placed on a single bench in a walk-in growth chamber set to 10-hour light and 14-hour dark cycles. Daytime temperature was maintained at 24°C, while nighttime temperature was kept at 20°C.

### 3.6 Harvesting and Phenotyping

Plants were harvested under a laminar flow hood after 31 days in the growth chamber. The roots were cut using autoclaved scissors, flamed with ethanol between each sample. The roots were rinsed with epiphyte buffer to remove clay particles, dried on individual paper towels that had been autoclaved, and placed onto pre-cut sheets of aluminum foil. The epiphyte removal buffer was prepared by dissolving 3.75 g KH2PO4, 4.75 g K2HPO4, and 0.5 ml Triton X-100 (Fisher Scientific, Waltham, MA, USA; CatNo. NC0481054) in deionized water to a final volume of 500 ml, followed by filtration through a 0.2 µm filter to sterilize the solution. The foil was folded, labeled, and placed into liquid nitrogen immediately, and samples were subsequently stored in −80C until processing. The aboveground biomass from each microbox (all three plants together) was placed in individual envelopes. The envelopes were dried in an oven at 60°C for 3 days and the dried plant samples were weighed to determine their dry weight.

### 3.7 Root sample processing

Forceps sterilized with bleach, ethanol, and flamed between each sample were used to remove the roots from the aluminum foil to be ground via mortar and pestle. Each sample was processed using a separate, autoclaved mortar and pestle, which was cooled using liquid nitrogen prior to grinding. After grinding, each individual sample was stored in sterile 15 mL Falcon tubes chilled with dry ice. The tubes were subsequently stored at −80°C until DNA extraction.

### 3.8 DNA extraction and 16S rRNA library preparation

DNA extraction was performed using the PowerSoil Pro Kit from DNeasy following the manufacturer’s instructions. The extracted DNA was eluted in RNAse-free water and stored at −80 C.

### 3.9 Amplicon sequence processing and analysis

We quantified extracted DNA (Invitrogen, Waltham, MA, USA; Quant-iT dsDNA Assay Kit, CatNo. Q33120) and normalized all samples to equal concentrations prior to PCR. All PCRs were performed in duplicate, each reaction consisting of 15 uL DNA template at 1 ng/uL, 18.75 uL PlatinumII Hot-Start PCR Master Mix (2x) plus 0.75 uL BSA (20 mg/mL), 0.285 uL of 100uM mitochondrial PNA (GGCAAGTGTTCTTCGGA; PNA Bio, Thousand Oaks, CA, USA; CatNo. MP01-50), 0.285 uL of 100uM chloroplast PNA (GGCTCAACCCTGGACAG; PNA Bio, Thousand Oaks, CA, USA; CatNo. PP01-50), 0.75 uL of 10uM of each forward and reverse primer targeting the V3-V4 region of the 16S rRNA gene, and water to bring final volume to 37.5 uL. From 5’-3’ complete primer sequences included an Illumina adapter sequence, a 12 base pair index, an Illumina TruSeq primer sequence, a 1-7 base pair spacer to increase flow cell diversity, followed by forward (CCTACGGGNBGCASCAG) and reverse (GACTACNVGGGTATCTAATCC) 16S rRNA primer sequences; a complete list of primer sequences is available in Supplemental Table 2. Thermocycling conditions were set to follow: an initial 3 min cycle at 94°C, then 36 cycles of 1 min at 94°C, 30 sec at 48°C, 1 min at 72°C, and a final cycle of 10 min at 72°C before holding at 4°C indefinitely. Replicate PCRs were then pooled and quantified before pooling all libraries in equimolar amounts. Pooled libraries were then cleaned with AMPure XP beads (Beckman Coulter, Brea, CA, USA; CatNo. A63880) following the manufacturer’s protocol, eluted in water, then submitted to University of California, Berkeley’s QB3 Genomics for Illumina MiSeq v3 300bp paired-end sequencing (Berkeley, CA, RRID:SCR_022170; Illumina, San Diego, CA, USA).

### 3.10 Sequencing analysis

Demultiplexed reads were imported into QIIME2 (Bolyen et al. 2019) where primer sequences were removed using the Cutadapt plugin (Marcel 2011). Reads were then trimmed to remove low-quality base pair calls before being passed to the DADA2 plugin (Callahan et al. 2016) for denoising and dereplication, resolving sequences into amplicon sequence variants (ASVs).

Representative sequences of ASVs were then assigned taxonomies via the feature-classifier plugin (Bokulich et al. 2018) using the sklearn method (Pedregosa et al. 2012) and a classifier trained on the SILVA 138 SSU Ref NR 99 16s rRNA reference database (Quast et al. 2013). Prior to downstream analyses, we filtered ASVs identified as either mitochondria or chloroplast.

The corresponding data tables of ASV frequency and taxonomy were manipulated using a variety of packages including dplyr (Wickham et al. 2023), tidyr (Wickham, Vaughan, and Girlich 2024), and tibble (Wickham 2016b). Diversity metrics and NMDS scores were calculated using the vegan R package (Oksanen et al. 2024)while differential abundances were calculated using ANCOM-BC2 (Lin and Peddada 2024). Finally, the data was plotted using ggplot2 (Wickham 2016a) and pheatmap (Kolde 2018).

## DISCUSSION

### 4.1 Conserved plant growth promotion

In this study we observed dramatically increased biomass yield when plants were grown in the presence of a living microbiome and organic matter (OM). Since the plants were grown under nutrient limiting conditions and we did not observe growth promotion by living microbiomes without OM, the increased yield was most likely due to the release of plant-available nutrients from the decomposition of OM. Interestingly, microbiomes from diverse soils or unautoclaved OM uniformly increased plant biomass despite stark differences in their bacterial community composition. It should be noted that the leaves harvested for the OM used in this study carried a microbiome that may have contained a variety of epiphytic microbes as well as soil microbes due to its small stature adjacent to the soil surface. This suggests that each microbiome used in this study possessed the functional capacity to break down organic matter and supply essential nutrients to the plants. A variety of studies have demonstrated that microbiome origin impacts microbiome composition without necessarily altering plant growth phenotypes (Pérez-Jaramillo et al. 2017; Brown et al. 2020; Liu et al. 2019). Several studies have demonstrated that despite varied taxonomic composition, soil microbiomes largely retain the same core functions (Louca, Parfrey, and Doebeli 2016), though the frequency of genes impacting certain processes can vary between microbiomes adapted to different conditions, including plant decomposition traits (Fierer et al. 2012). It is plausible that soil microbial communities that rely on plants for their carbon sources, such as those used in this study, all face selective pressures to maintain the ability to efficiently decompose plant material. The OM used in this study, green grass leaves, is readily digestible so perhaps it is not surprising that all the microbiomes were able to degrade it equally. A significant effect of microbiome origin may have been obtained if a more recalcitrant form of OM (e.g. wood, pine needles) was used.

However, even in divergent microbial communities, the ability to decompose complex organic substrates and release nutrients to support plant growth is highly conserved. This is in contrast to studies that have examined other stressors, such as salt stress and drought, which showed differences in soil origin significantly impacted plant phenotype in response to these stressors (Lau and Lennon 2012; Santos et al. 2021). The association of similar phenotypes despite varying 16S data suggests that while these microbiomes may differ taxonomically, their metabolic capabilities related to decomposition and nutrient acquisition by living plants remain intact due to high selective pressure to retain these traits.

### 4.2 Experimental system provides clear selection for microbes related to decomposition

The experimental system used in this study was created in part with the goal of creating a simplified microbiome that performs functions related to plant growth promotion via decomposition from an initially complex mixture of soil microbes. This could be performed by creating SynComs via a variety of methods, such as isolating microbes that persist after successive passaging of microbiomes in this system. This strategy has been utilized in other studies to create stable, less-diverse microbial communities that show apparent habitat-filtering, in which a specific condition selects for a subset of microbes (Morella et al. 2020).

Environmental stressors, such as drought (Naylor et al. 2017; Lauber et al. 2009)), pH (Lauber et al. 2009), and unavailability of organic carbon sources (Lauber et al. 2009; Cesarano et al. 2017; Testen and Miller 2018), have also been identified to decrease microbial diversity. In the present study, the variability of retained diversity and species richness across different soil inoculum treatments suggests that each may have had a different capacity to be mined for microbial genetic resources targeting decomposition. Additionally, specific differences in alpha diversity and species richness between treatments were informative. For example, the experimental samples given no organic substrates (unplanted, no organic matter supplementation) had no detectable 16S sequences at the end of the experiment, demonstrating the requirement for external carbon to allow bacterial growth to detectable levels, as the calcined clay substrate is completely inorganic. Additionally, no or low contamination of the controls suggest the different microbiomes were effectively isolated from one another during the course of the experiment. Furthermore, the compost inoculum, which has by definition undergone selective successive passaging, was shown to better retain its diversity and species richness compared to the inoculum from soils. Collectively, these data show that the experimental design was effective in reducing the community to subsets of microbes that could persist in the specific conditions of the experiment, and that utilization of this experimental technique allows for effective targeting of microbes or microbial communities that provide functions related to decomposition.

### 4.3 Specific microbial taxa implicated in plant growth promotion and decomposition

The experimental system described in this study was effective at selectively reducing the microbial diversity from the inoculum, while simultaneously resulting in a community of microbes that maintain their ability to perform decomposition. The identification of conserved functional niches for specific bacterial lineages across multiple ecosystems is quite common; microbiomes from entirely different continents have been found to have a phylogenetically conserved response to soil nitrogen addition (Isobe et al. 2019) as well as simulated perturbations (Martiny, Treseder, and Pusch 2013). In this study, several such lineages were identified, including the genera Achromobacter and Caulobacter. Both of these genera were enriched under OM treatments in multiple soil treatments, and have been found in other studies to contain strains that have plant growth promotive properties(Jha and Kumar 2009; Abdel-Rahman et al. 2017; Ma, Rajkumar, and Freitas 2009; Luo et al. 2019; Berrios 2021; Berrios and Ely 2020). The most striking example is the genus *Massilia* due to its conserved presence, high prevalence, and specific pattern of differential abundance (Figure 4B, S1-2) which suggest it likely has a conserved functional role in the decomposition process. Species in the genus *Massilia* are often described as plant associated, though they have been identified and isolated from a wide range of diverse and often extreme environments (Baek et al. 2022; C. Li et al. 2021; Manni and Filali-Maltouf 2022; Sedláček et al. 2022). Some *Massilia* species have been shown to promote plant growth (Guo et al. 2019) and pathogen resistance (C. Li et al. 2021). Perhaps most relevant for this study, bacteria within this genus isolated from the rice rhizosphere have been shown to degrade cellulose (Du et al. 2021) and other compounds found in organic matter (Sedláček et al. 2022). We suspect that the role of *Massilia* and other decomposition-related bacteria identified in this study likely beneficially impact plant phenotype through their decomposition functions. Alternatively, their correlations with OM treatment may be the result of interactions with other microbial taxa and a byproduct of other community dynamics without affecting plant phenotype. The same taxa sometimes respond to the same changes in soil conditions despite being on different continents, suggesting that some within the genus might have a conserved function (Martiny, Treseder, and Pusch 2013). Further analysis of the bacteria within the *Massilia* genus will allow for more insight into which (if any) mechanisms this group of bacteria utilizes to promote plant growth and whether its clear association with decomposition is causal.

### 4.4 Conclusions and future directions

Through careful use of controls, we showed that decomposition of plant organic matter by the microbiome, no matter what soil that microbiome is adapted to, promotes plant growth. The conserved function of ecologically distinct soil microbiomes in promoting the early growth of seedlings of a plant that evolved on a separate continent implies an ancient adaptation of soil microbiomes to promote plant growth in a mutualism resulting from the need of soils to consume plant derived carbon. This experimental system provides the means by which to begin teasing which functional consortia might be responsible for plant growth promotion due to decomposition. Microbiome analyses in this experiment point to the persistence of this function across both a wide taxonomic breadth as well as potentially within specific taxa, such as the Massilia genus, which would be of particular interest for further studies. Additionally, owing to its robust reproducibility, the system could be utilized to further characterize and validate the function of microbes identified to be associated with plant growth promotion via isolation and monoassocation assays, syncoms, metagenomics, trancriptomics, or any combination thereof. Characterization of such mechanisms will allow for microbes related to plant growth promotion via decomposition to be studied thoroughly in an effort to potentially increase plant yield in agricultural settings by creating favorable conditions for nutrient acquisition from OM by plants via microbes.

## Funding

The work conducted by the U.S. Department of Energy Joint Genome Institute (https://ror.org/04xm1d337), a DOE Office of Science User Facility, was supported by the Office of Science of the U.S. Department of Energy operated under contract no. DE-AC02-05CH11231, and the US Department of Agriculture under CRIS project no. 2030-12210-003-000D.

## Declaration of generative AI and AI-assisted technologies in the writing process

During the preparation of this work the author(s) used ChatGPT to aid in content structuring. After using this tool/service, the author(s) reviewed and edited the content as needed and take(s) full responsibility for the content of the published article.

## Data availability

Data can be found on the NCBI Sequence Read Archive via project identifier PRJNA1161380.

## CRediT authorship contribution statement

Bradie Lee: Conceptualization, Data curation, Formal analysis, Investigation, Methodology, Validation, Visualization, Writing - original draft. Devin Coleman-Derr: Funding acquisition, Project administration, Resources, Supervision, Writing - review and editing. John P. Vogel: Funding acquisition, Project administration, Resources, Supervision, Writing - review and editing.

## Acknowledgements

Thank you to Mingqin (Mike) Shao for his assistance in harvesting the experiments. Thank you to the members of the Vogel and Coleman-Derr labs for their continual feedback and critique during the development of the experimental system.

## Declaration of competing interest

The authors declare that they have no known competing financial interests or personal relationships influencing the work presented here.

---

**Table S1).**
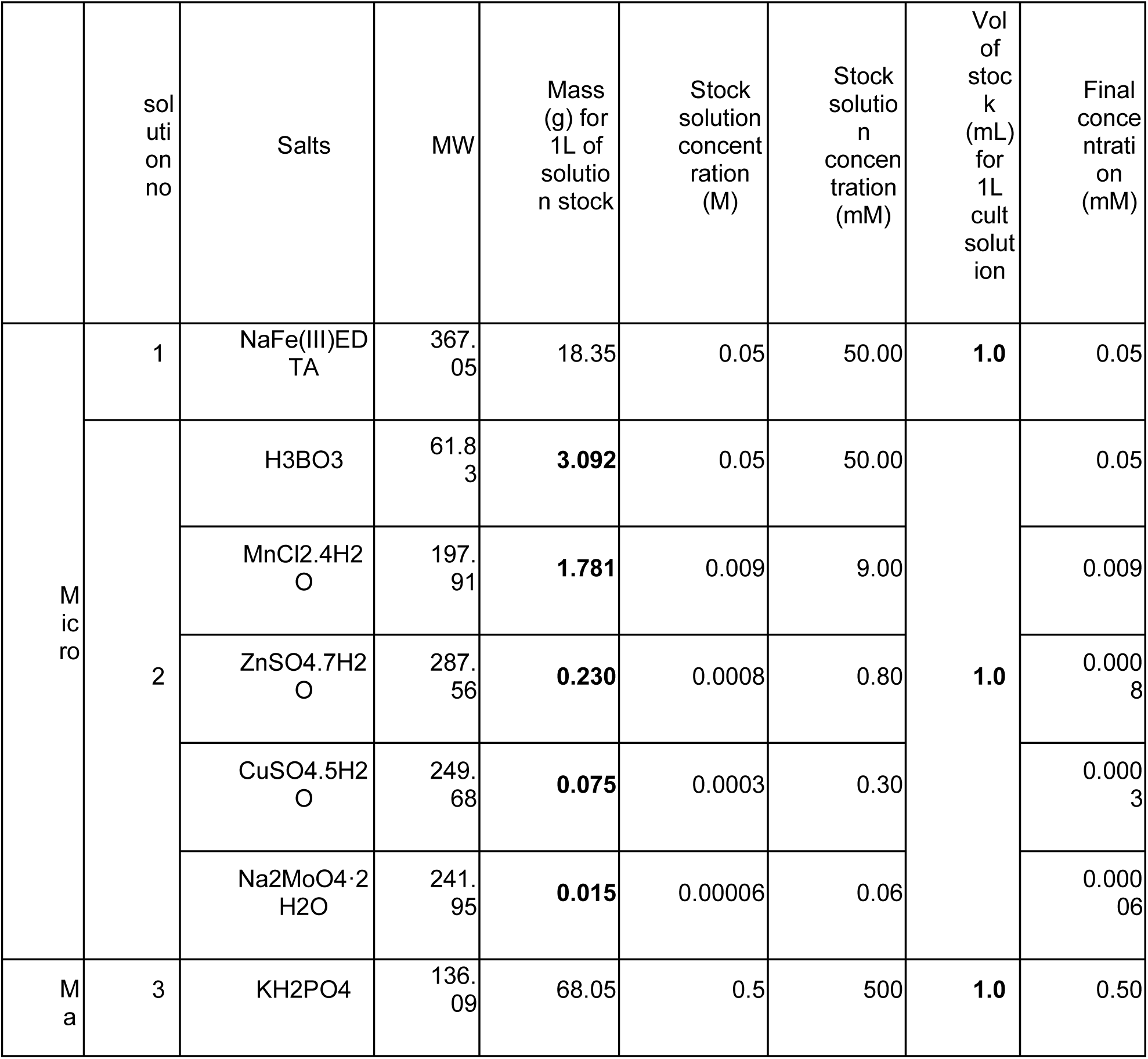
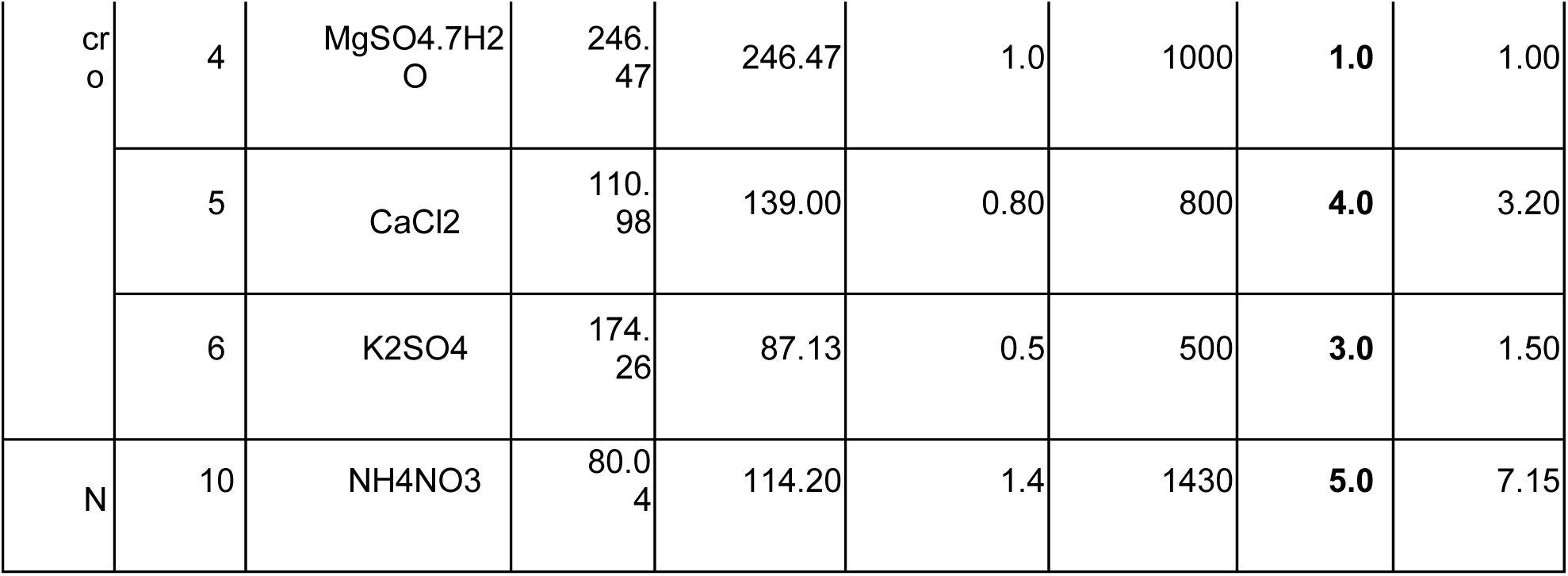

**Figure S1).**
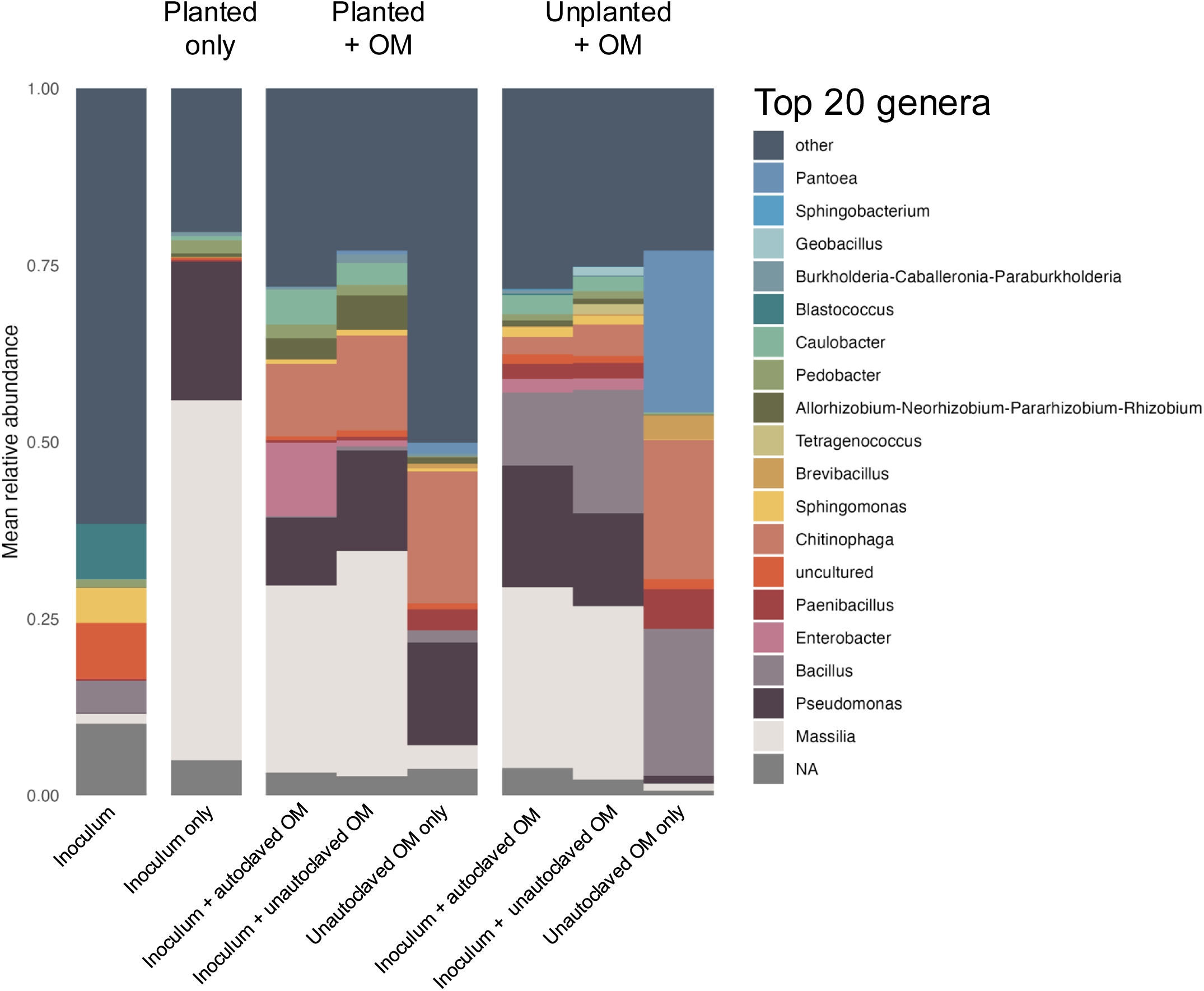
Relative abundance barplot displaying the top 20 genera present in the samples that contained a living microbiome. Each bar shows the mean for all samples within each treatment type.

**Figure S2).**
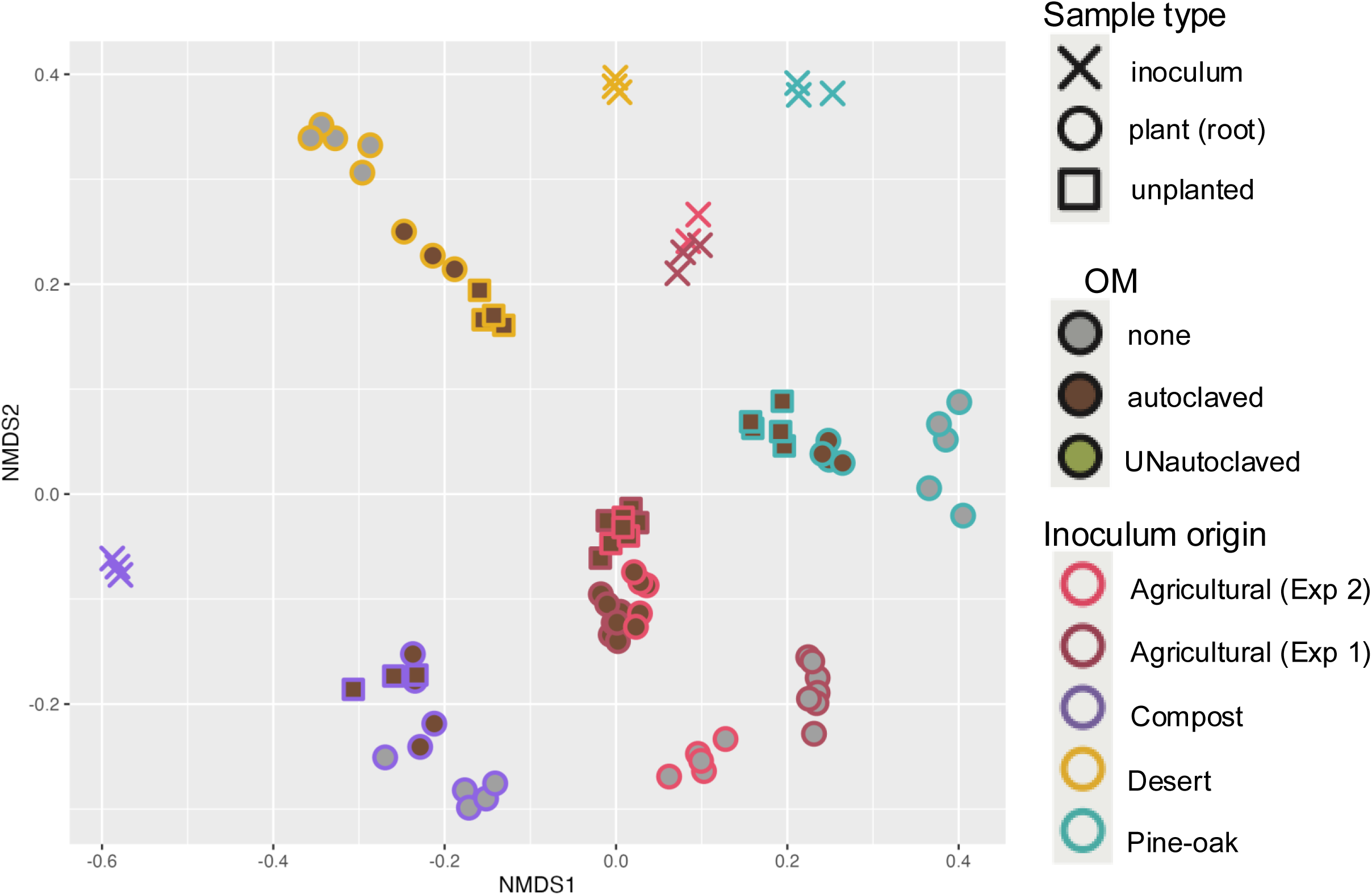
Bray-Curtis NMDS plot displaying samples containing living microbiomes in the second experiment as well as analogous samples in the first experiment

**Figure S3.**
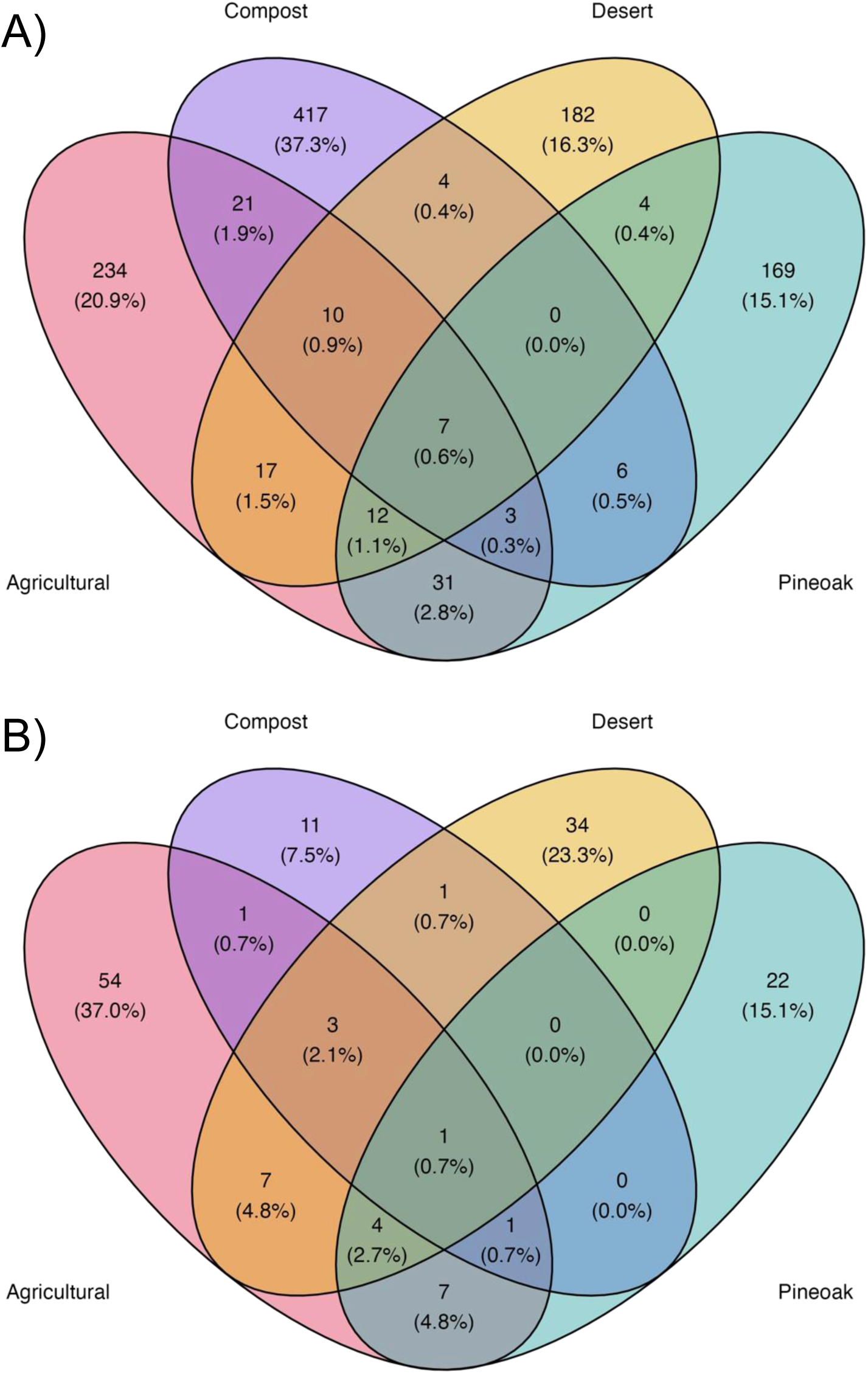
A) Venn diagram showing overlap of individual ASVs found in Experiment B with over 100 reads per ASV per inoculation type. B) Venn diagram showing overlap of individual ASVs classified as Massilia.

**Figure S4.**
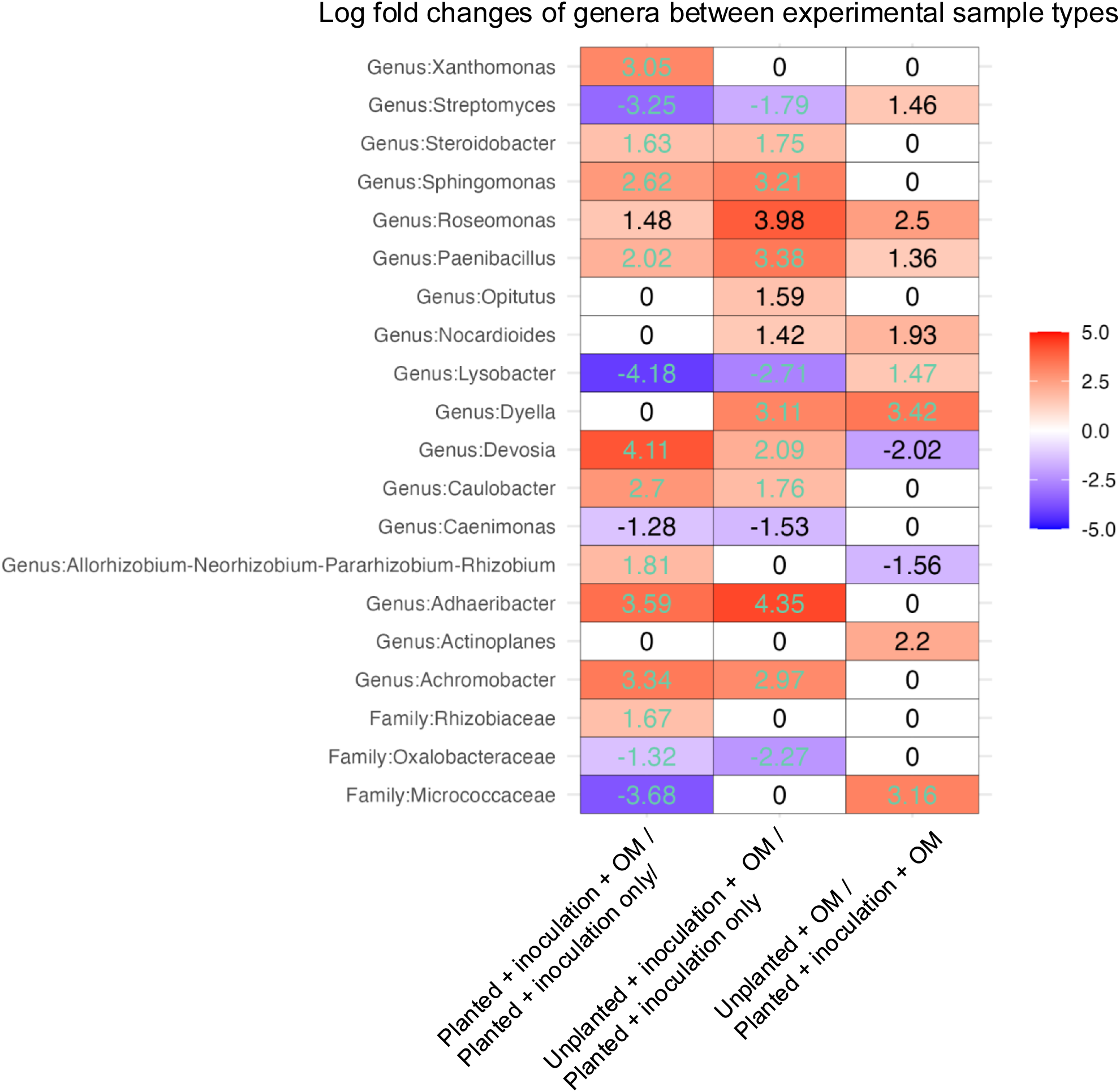
Log fold changes of taxa that were found to be significantly differentially abundant genera across samples from experiment A, excluding inoculum samples. Teal numbers represent those comparisons that passed the ANCOM-BC2 sensitivity analysis, black did not.

**Figure S5.**
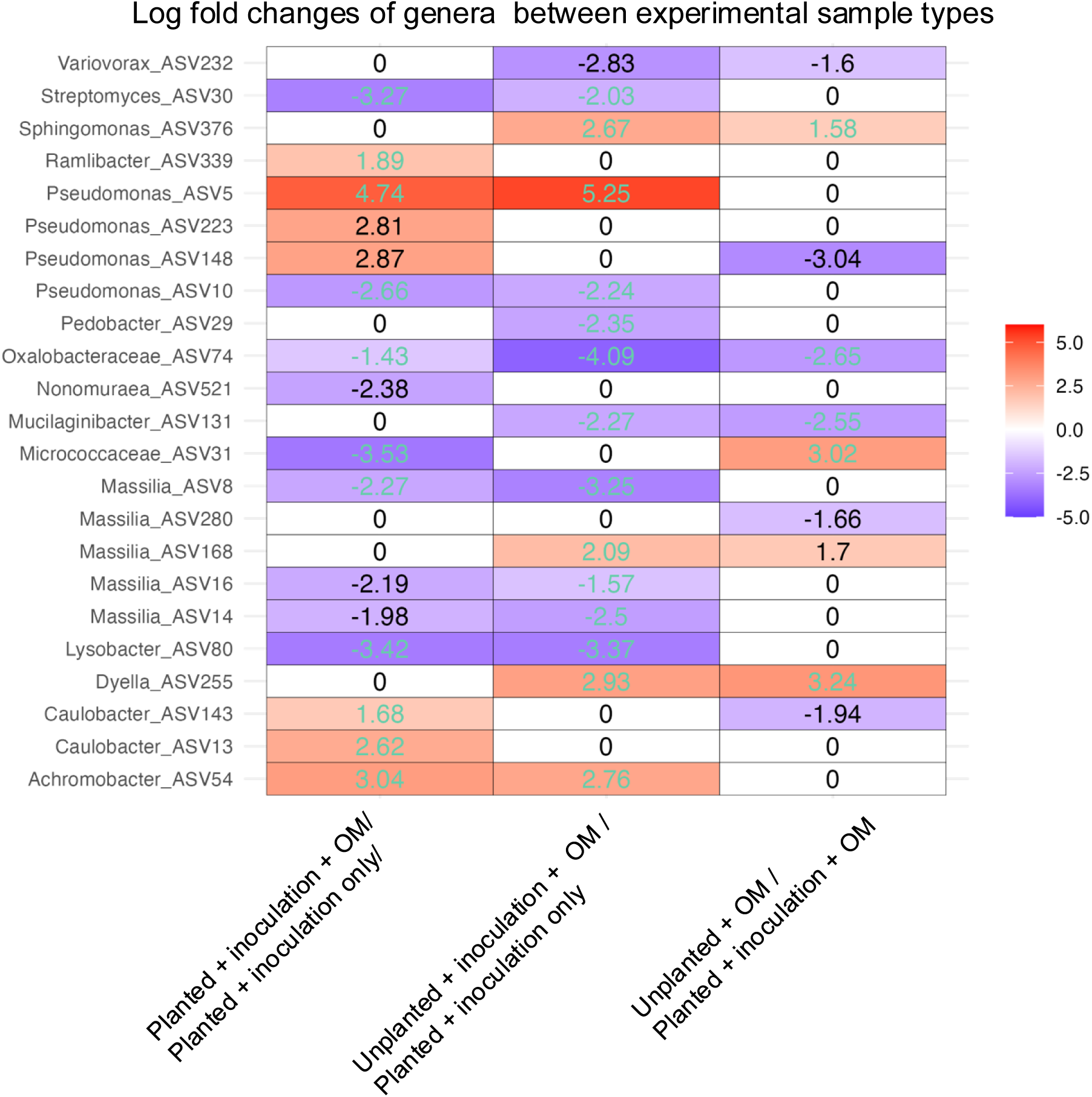
Log fold changes of ASVs that were found to be significantly differentially abundant when utilizing ANCOMBC2 across samples from experiment A, excluding inoculum samples. Teal numbers represent those comparisons that passed the ANCOM-BC2 sensitivity analysis, black did not.

**Figure S6.**
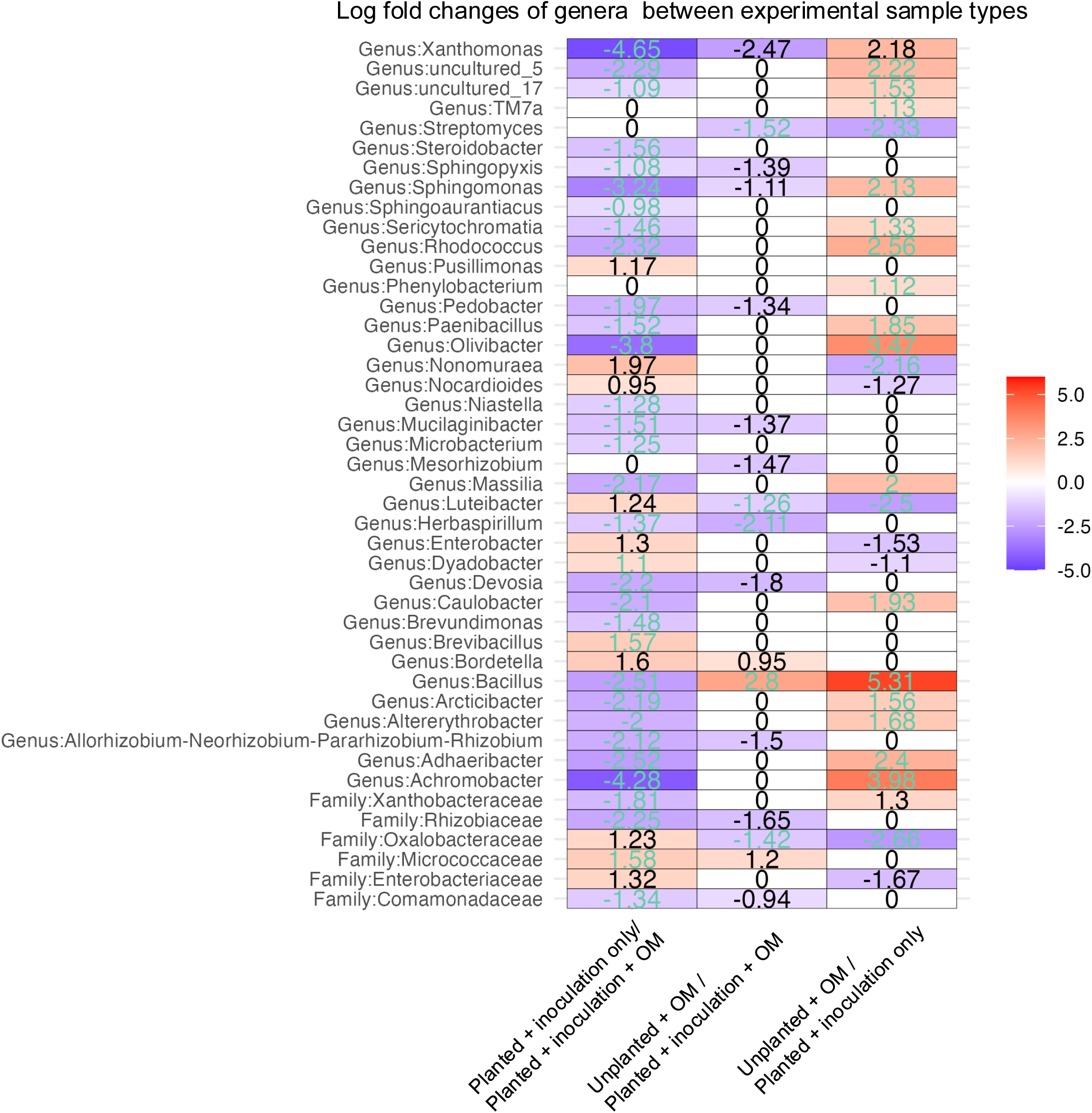
Log fold changes of genera that were found to be significantly differentially abundant when utilizing ANCOMBC2 between inoculation types from experiment B, excluding inoculum samples. Teal numbers represent those comparisons that passed the ANCOM-BC2 sensitivity analysis, black did not.

**Figure S7.**
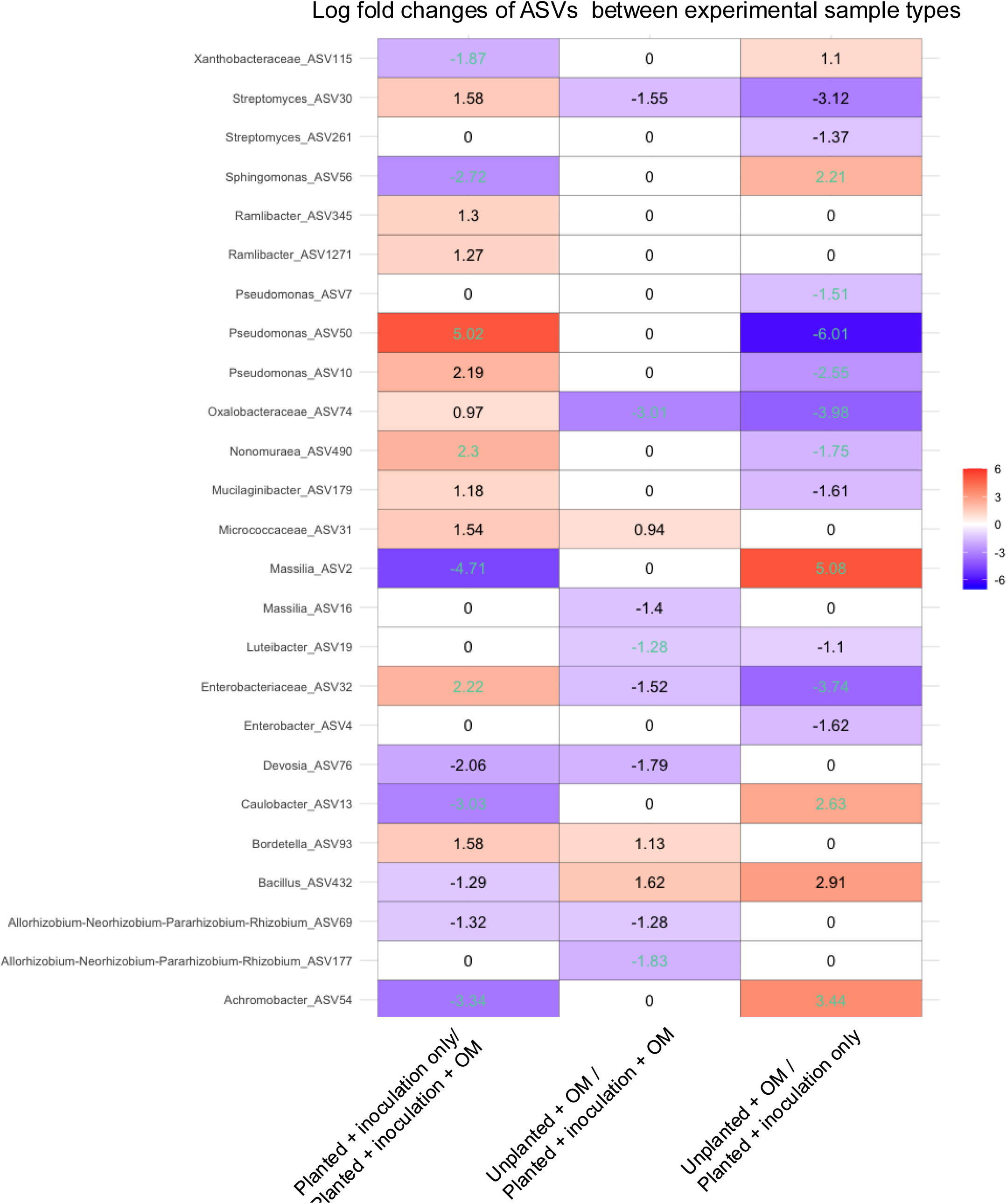
Log fold changes of ASVs that were found to be significantly differentially abundant when utilizing ANCOMBC2 between inoculation types from experiment B, excluding inoculum samples. Teal numbers represent those comparisons that passed the ANCOM-BC2 sensitivity analysis, black did not.

**Figure S8.**
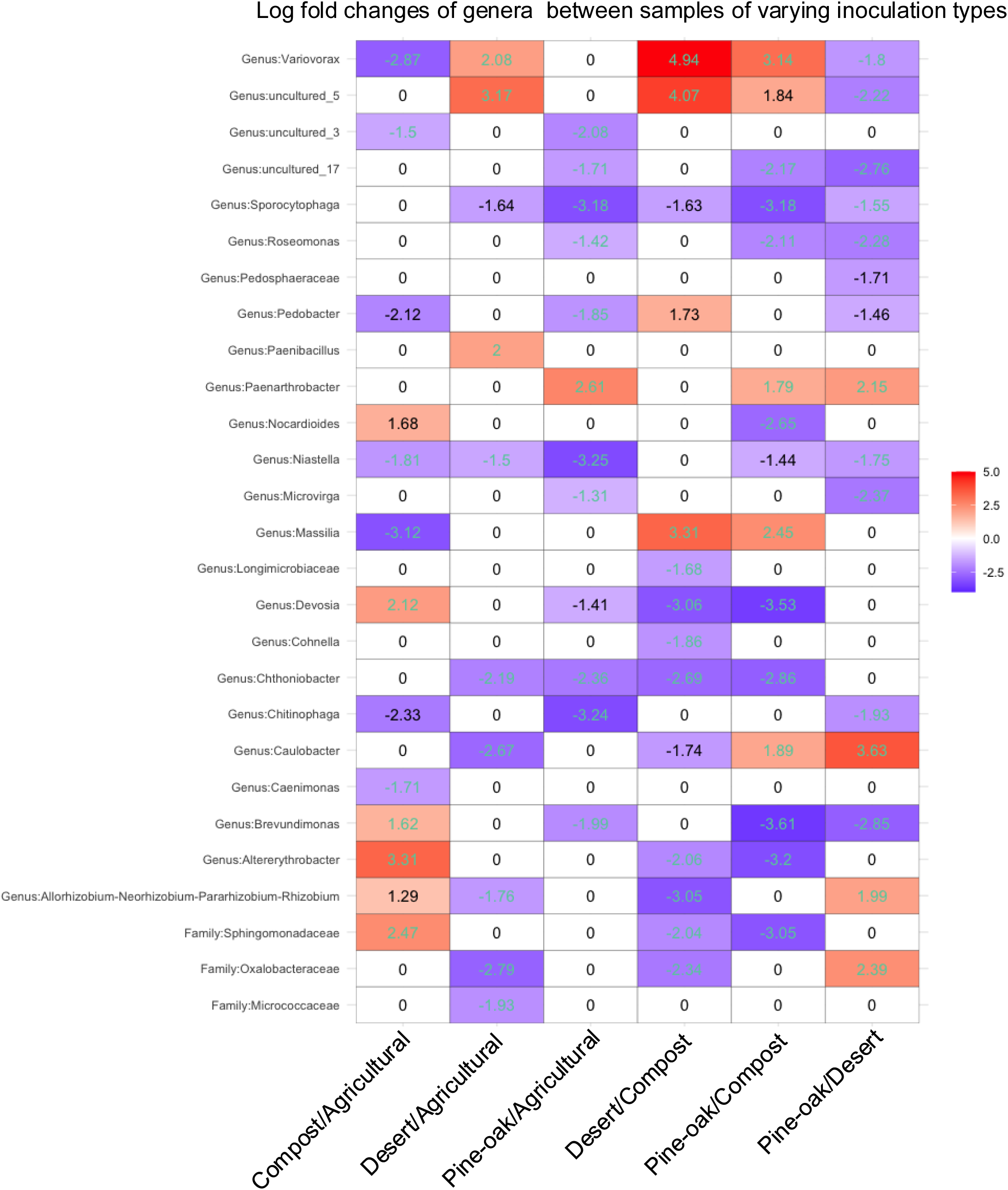
Log fold changes of genera that were found to be significantly differentially abundant when utilizing ANCOMBC2 between inoculation types from experiment B, excluding inoculum samples. Teal numbers represent those comparisons that passed the ANCOM-BC2 sensitivity analysis, black did not.

**Figure S9.**
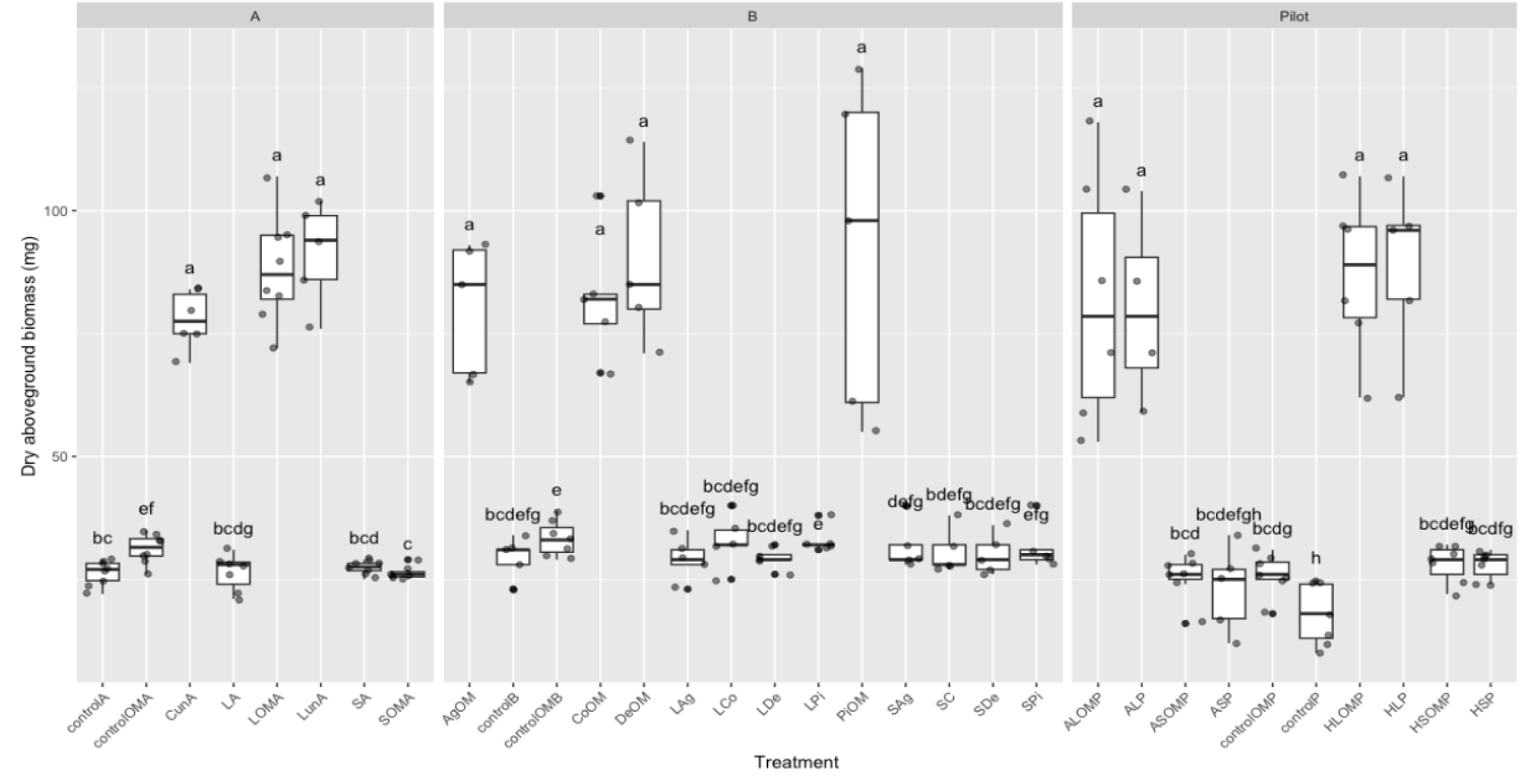
Phenotype differences in the first, second, and a pilot experiment utilizing the same experimental system and protocol. Letters indicate pairwise comparison differences between treatments. All bars labeled “a” represent treatments that received a combination of a living microbiome and OM, while all others were some combination of other factors.

## Notes

### Competing Interest Statement

The authors have declared no competing interest.

